# Applying Machine Learning to Classify the Origins of Gene Duplications

**DOI:** 10.1101/2021.08.12.456144

**Authors:** Michael T.W. McKibben, Michael S. Barker

## Abstract

Nearly all lineages of land plants have experienced at least one whole genome duplication (WGD) in their history. The legacy of these ancient WGDs is still observable in the diploidized genomes of extant plants. Genes originating from WGD—paleologs—can be maintained in diploidized genomes for millions of years. These paleologs have the potential to shape plant evolution through sub- and neofunctionalization, increased genetic diversity, and reciprocal gene loss among lineages. Current methods for classifying paleologs often rely on only a subset of potential genomic features, have varying levels of accuracy, and often require significant data and/or computational time. Here we developed a supervised machine learning approach to classify paleologs from a target WGD in diploidized genomes across a broad range of different duplication histories. We collected empirical data on syntenic block sizes and other genomic features from 27 plant species each with a different history of paleopolyploidy. Features from these genomes were used to develop simulations of syntenic blocks and paleologs to train a gradient boosted decision tree. Using this approach, Frackify (Fractionation Classify), we were able to accurately identify and classify paleologs across a broad range of parameter space, including cases with multiple overlapping WGDs. We then compared Frackify with other paleolog inference approaches in six species with paleotetraploid and paleohexaploid ancestries. Frackify provides a way to combine multiple genomic features to quickly classify paleologs while providing a high degree of consistency with existing approaches.

## 1. Introduction

Polyploidy, or whole genome duplication (WGD), is a common phenomenon among some lineages of eukaryotes, particularly flowering plants where nearly 30% of extant species are estimated to be recent polyploids ***(1–3)***. Although many of these polyploid species ultimately go extinct ***(4–7)*** many return to the diploid state through a process known as diploidization ***(8)***. This is a ubiquitous process as most land plants are ancient polyploids that have diploidized, with several lineages containing multiple WGD in their recent and ancient history ***(9)***. WGD derived genes, also known as paleologs or ohnologs, are rapidly lost during diploidization in a process termed fractionation ***(8, 10–12)***. The paleologs that are not lost during this process can persist in diploidized genomes for millions of years ***(13, 14)***. Paleologs are thought to play important roles in a variety of evolutionary processes including speciation through reciprocal gene loss or lineage specific rediploidizaiton ***(15–21)*** as well as adaptation and the evolution of novelty by sub- and neofunctionalization ***(22–26)***. Recent analysis has shown that paleologs have higher levels of genetic diversity than non-paleologs and can significantly contribute to domestication ***(14)***. Further exploration into the legacy of ancient polyploidy requires the rapid and accurate identification of paleologs in the genomes of extant diploid descendants.

Current methods for the identification of paleologs utilize sequence comparisons, genome structure information, or a combination of the two. Arguably the most common and simplest method used to detect paleologs is to identify peaks of gene duplication by examining gene age distributions ***(27–31)***. WGD instantly doubles the number of genes in a genome and this large, abrupt birth event is visible in gene age distributions. These age distributions are often called Ks plots (Fig. 1) because they are usually made by plotting the synonymous site divergences (Ks) among paralogs in a single genome. Mixture models can be used to identify and place bounds on peaks of gene duplication in Ks plots, and homologous sets of genes within these Ks bounds are assumed to be paleologs ***(27–29)***. Although simple and applicable to a wide range of data, this approach suffers from potentially high rates of false positives and negatives. For example, not all genes within a Ks peak were necessarily duplicated by a WGD and the hard bounds placed by mixture models are not perfectly accurate (Fig. 2). Another method for identifying paleologs is to leverage genome structure rather than only comparing unordered sequences. This approach uses conserved gene order—synteny—either within a species or between an outgroup to identify the duplicated regions in genomes from WGDs ***(32–36)***. Syntenic regions can also arise from other small scale duplications or additional layers of older WGD ***(23, 37, 38)***.

**Figure 1:**
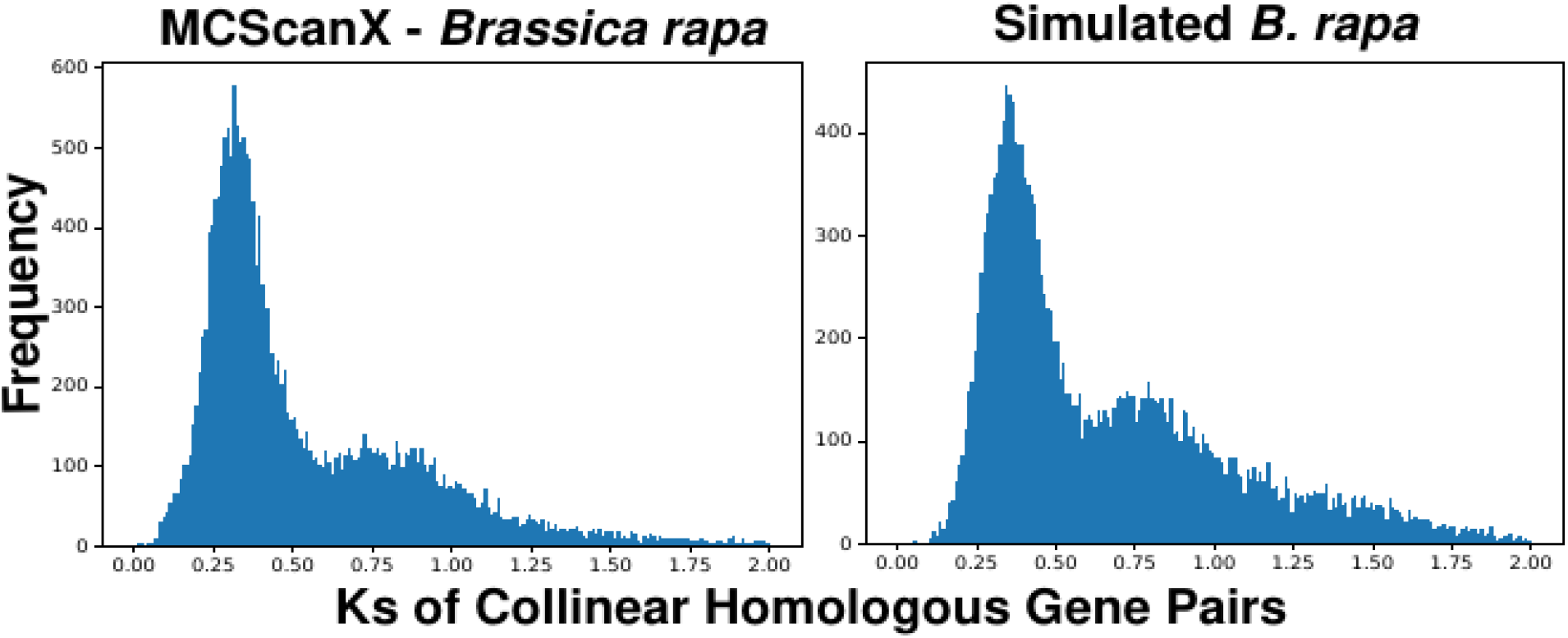
Comparison of the distribution of collinear homologous gene pair Ks_pair_ from syntenic block pairs in MCScanX results from *Brassica rapa* vs simulated MCScanX results.

**Figure 2:**
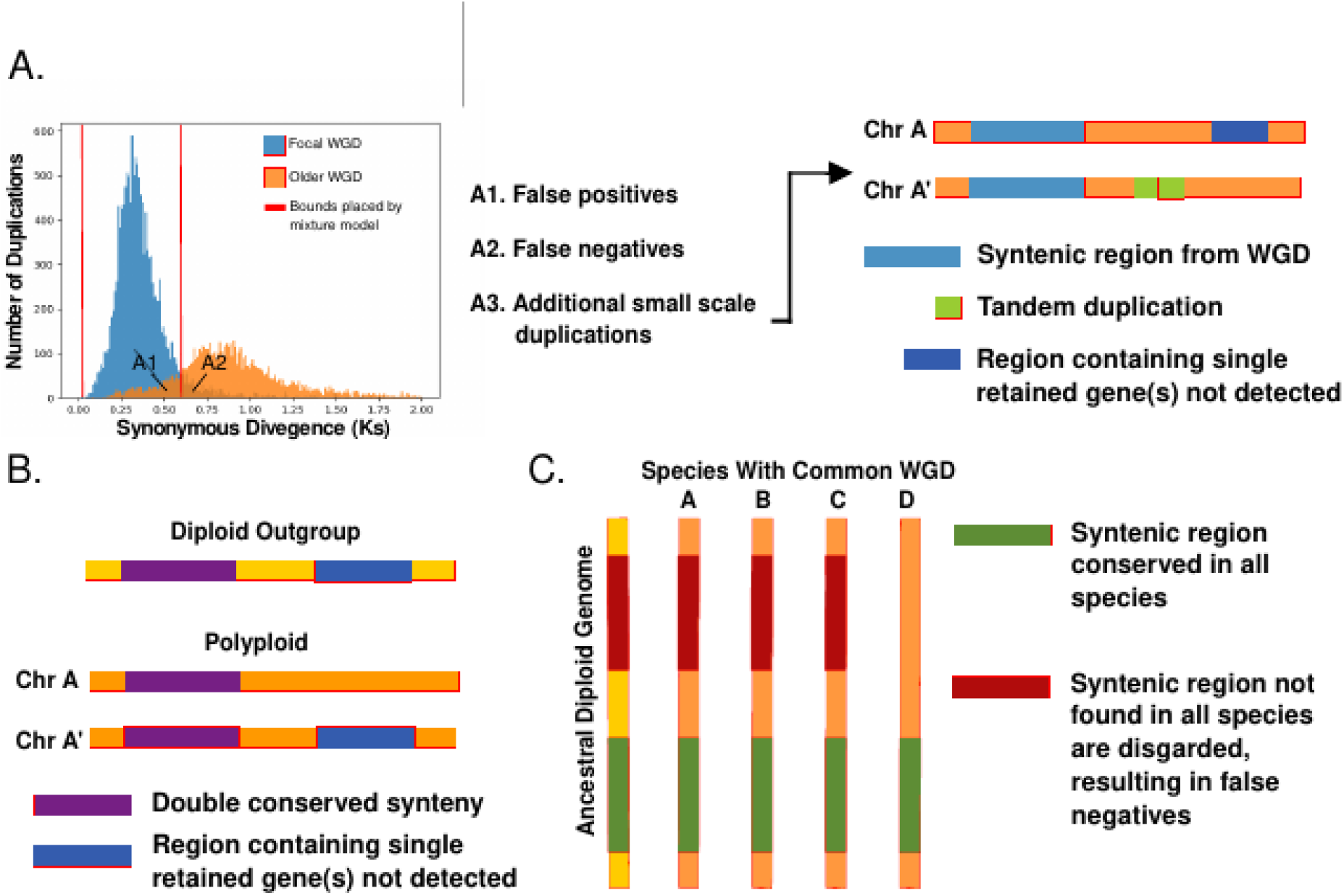
Examples of major sources for false positives and negatives in a variety of paleolog identification approaches. A.) Hard bounds placed by mixture models can produce false positives and negatives (A1, A2) while also mis-identifying paralogs from small scale duplications as paleologs. By using syntenic information some paralogs from small scale duplications can be excluded at the cost of missing single retained paleologs (A3). B.) Using DCS blocks improves identification of paleologs that are still present in two or more copies, but can lead to exclusion of single retained paleologs that are missing their other paralog(s). C.) Approaches such as POInT discards regions that are not conserved across all species used in the analysis, resulting in false negatives.

Several methods have been employed to resolve the origins of syntenic regions. The simplest of these approaches is to compare the overlap of genes within syntenic regions to that of the Ks based methods ***(14, 34, 39–41)***. As an alternative to Ks, some studies have used Double Conserved Synteny (DCS) with a diploid outgroup that does not share the WGD. In DCS analyses, syntenic regions in the self-self comparison that match to the homologous syntenic block in the outgroup are considered to be WGD duplicates (Fig. 2)***(35, 36)***. Although these methods can be useful for filtering out small scale duplications, both suffer when multiple WGDs are present, a common occurrence in plants ***(9)***. Relying solely on syntenic information will result in DCS being detected not only for the focal WGD, but possibly also older WGD ***(34, 42)***. Filtering these blocks by Ks can also be insufficient because the overlap between adjacent WGDs can make them difficult or impossible to separate (Fig. 2)***(34, 43)***.

To help disentangle the origins of syntenic genes, several methods have been developed that account for the complex duplication histories of plants. One approach, quota-align, attempts to differentiate paralogous and orthologous syntenic blocks by using Binary Integer Programming (BIP) to fit syntenic blocks that match to an *a priori* chosen ploidy level ***(34)***. Although this method can be useful for disentangling the origin of individual syntenic blocks, it often fails to identify genes retained in the single copy state as they are inherently undetected due to missing their homolog (Fig. 2). This method is also limited by the user’s *a priori* knowledge of the evolutionary history of their study organism ***(34)***. As an alternative, POInT (Polyploid Orthology Inference Tool) provides a powerful framework for reconstructing the evolutionary history of ancient WGDs, and is one of the few methods to formally model WGD evolution Although paleolog classification is not the primary goal of POInT, its inferences can be used to identify paleologs ***(11, 44)***. An inherent drawback to POInT’s approach is that inferences are only made for regions with conserved synteny across all species examined, resulting in many paleologs being missed, especially those in potentially interesting areas of genomes that have diverged in retention or experienced high rates of rearrangement since a WGD ***(11, 44)***. For sets of paleologs that are detected, the retention status of each copy can be filtered by the model confidence, limiting error in downstream analysis ***(11, 44)***. The tradeoff of these sophisticated models is that it requires non-trivial computational effort to prepare data and run POInT; including the identification of high confidence DCS blocks within species using simulated annealing and the estimation of ancestral gene order using DCS blocks conserved across all species of interest. Overall, each of the existing approaches to identify paleologs have strengths and weaknesses that can be leveraged to provide a specialized tool for paleolog identification and classification that is computationally efficient.

Many large scale and complex genomic questions have eluded evolutionary biologists due to computational limitations or gaps in theory. Machine learning based approaches have recently been used to overcome some of these limitations, especially in cases where the biology can be simulated ***(45)***. Recent examples include the detection of selective sweeps ***(46, 47)***, describing past demographic history and recombination ***(48–51)***, landscape connectivity ***(52)***, inferring phylogenetic structure ***(53)***, species delimitation ***(54)***, and hybrid speciation and admixture ***(55)***. The benefit of supervised machine learning in these cases is that they often achieve accurate inferences with much less computational time and effort than traditional inference methods ***(53, 56)***. In the context of identifying paleologs, machine learning allows for an ensemble approach that can leverage the strengths of various methods such as quota-align, DCS, and Ks divergence, yet also allows for the easy integration of updated methods in the future. Here we introduce a gradient boosted classification approach to identify paleologs using multiple features from synteny and Ks information. This method, Frackify (Fractionation-Classify), is able to make inferences with a high degree of overlap with POInT classifications in conserved syntenic regions while also identifying putative paleologs in regions not analyzed by POInT. We begin with exploring biological data in order to make informed simulations and the caveats of such an approach. We then explore methods for training, testing, and validation a gradient boosted decision tree. Finally, we analyze six paleohexaploid and paleotetraploid plant genomes to compare Frackify with other paleolog inference approaches.

## 2. Materials

### 2.1 Dependencies

Frackify is a python 3 based pipeline that requires the following python libraries: numpy (1.18.1), matplotlib (3.1.3), sys, statistics, pandas (1.0.1), random, itertools, csv (1.0), os, scipy (1.4.1), xgboost (1.3.3), and pickle ***(57–63)***.

### 2.1 Pipeline Input

1. CDS file for focal species being analyzed
2. MCScanX or SynMap output with Ks/Kn information

a. Self - Self comparison
b. Self - Outgroup comparison
3. Mean Ks for each focal WGD in the species

Examples can be found in the Frackify GitLab repo: https://gitlab.com/barker-lab/frackify or Docker: https://hub.docker.com/r/mmckibben/frackify

## 3. Methods

### 3.1 Biologically Informed Approach

Frackify, like other machine learning based approaches, makes predictions using a series of “rules” which it infers from a training dataset where the “answers” are known ***(64, 65)***. Currently no such dataset exists for paleologs as methods for paleolog identification are not necessarily accurate. Insufficient training data is a common issue in bioinformatics and many researchers supplement training data using simulations ***(48, 55)***. However simulating WGD is often difficult or intractable given that many plant lineages have complex duplication histories ***(9, 11, 66–68)***. This issue is compounded by our incomplete understanding of genome evolution following WGD. A prime example is the ongoing exploration of how cytological and genetic diploidization interplay on evolutionary timescales ***(8)***. A more tractable approach is to simulate the detectable signatures of WGD in genomes. In our case we simulated the output of popular synteny inference tools such as MCScanX and CoGe SynMap ***(39, 69)***. Both of these tools produce output files containing inferred syntenic block pairs either within or between species. Additional settings also allow for Ks values to be produced for homologous gene pairs in the detected syntenic blocks ***(39, 69)***. To help inform our simulations using empirical datasets we used MCScanX to identify syntenic blocks in the genomes of 27 species (Table 1). These species were selected because they have well studied and assembled genomes across a variety of taxa, including 10 orders and 22 genera that have experienced varying numbers of paleotetraploidies or paleohexaploidies with different levels of divergence. Several key properties of these syntenic blocks were summarized and used to develop simulations of WGD to train Frackify.

**Table 1:** List of 27 species used for empirical analysis of syntenic block properties, including the version and source for each genome used in this study.

| Species | Genome Accessed | Source |
| --- | --- | --- |
| *Aethionema arabicum* | CoGe id23428 | (Nguyen et al. 2019) |
| *Ananas comosus* | CoGe id25735 | (Ming et al. 2015) |
| *Arabidopsis lyrata* | CoGe id3068 | (Rawat et al. 2015) |
| *Arabidopsis thaliana* | CoGe id1 | (Swarbreck et al. 2008) |
| *Brachypodium distachyon* | CoGe id25040 | (TIBI 2010) |
| *Brassica rapa* | CoGe id24668 | (X. Wang et al. 2011) |
| *Brassica juncea* | Brassica Database Bju15 | (Yang et al. 2016) |
| *Capsella rubella* | CoGe id16754 | (Slotte et al. 2013) |
| *Cleome violacea* | CoGe id23822 | (Emery et al. 2018) |
| *Cucumis melo* | MELONOMICS CM3.5.1 | (Testolin, Huang, and Ferguson 2016) |
| *Eutrema parvulum* | CoGe id12384 | (Dassanayake et al. 2011) |
| *Glycine max* | Phytozome 12 Wm82.a2.v1 | (Chang et al. 2013) |
| *Gossypium hirsutum* | NCBI Sequence Read Archive SRX202873 | (T. Zhang et al. 2015) |
| *Malus domestica* | Genome Database for Rosacea | (Jung et al. 2019) |
| *Oropetium thomaeum* | CoGe id25799 | (VanBuren et al. 2015) |
| *Oryza sativa* | Rice Genome Annotation Project V7.0 | (3,000 rice genomes project 2014) |
| *Phaseolus vulgaris* | Phytozome 12 version 1.0 | (Schmutz et al. 2014) |
| *Prunus persica* | Phytozome 12 version Prunus persica 2.1 | (Verde et al. 2013) |
| *Raphanus sativus* | Radish Genome Database RS1.0 | (H.-J. Yu et al. 2019) |
| *Ricinus communis* | Hale, http://castorbean.jcvi.org | http://castorbean.jcvi.org |
| *Setaria italica* | Phytozome 12 Setaria italica v2.2 | (Bennetzen et al. 2012) |
| *Sinapis alba* | CoGe id33284 | (G. Zhang et al. 2012) |
| *Solanum lycopersicum* | Phytozome 12 iTAG2.4 | (The Tomato Genome Consortium 2012) |
| *Solanum tuberosum* | Phytozome 12 Solanum tuberosum v4.03 | (Sharma et al. 2013) |
| *Sorghum bicolor* | Phytozome 12 Sorghum bicolor v3.1.1 | (McCormick et al. 2018) |
| *Thellungiella halophila* | CoGe id19492 | (Yang et al. 2013) |
| *Vitis vinifera* | Ensembl Plants release 51 | (Jaillon et al. 2007) |

A fundamental property of syntenic blocks is their length. In closely related taxa, syntenic blocks can be nearly as long as chromosomes, but over time they are broken up through fractionation and genomic rearrangements ***(8, 70, 71)***. To understand the lengths of syntenic blocks given the level of divergence of the WGD, we explored the distributions of self-self syntenic block lengths from MCScanX in our 27 exemplar species. Across the entire dataset, block lengths of less than 30 genes were predominant with a mean of 17.6 collinear gene pairs, however there was a long tail of much larger blocks with a maximum of 1285 genes in *Brassica juncea* (Fig 3). Although the approximate shape of a log-normal distribution was consistent across individual species, more fractionated genomes were skewed towards shorter block lengths. To test this, we fit a log-normal distribution to the distribution of syntenic block lengths in each species using the fitdistr() function from the MASS R library ***(72)***. We then plotted the mean and standard deviation of those distributions against the mean (Ks_WGD_) of collinearity homologous gene pair Ks (Ks_pair_) of the most recent WGD in each species (Fig. 3). Using a linear model we found that although the mean length of blocks did not vary significantly with Ks_WGD_ (F_1,27_= −1.909, p = 0.067, R^2^ = 0.123), it was a significant predictor of the standard deviation (Fig. 3)(F_1,27_= −2.242, p = 0.034, R^2^ = 0.162). Using this model we simulated the expected length of syntenic blocks for a given WGD by randomly selecting block lengths from a log normal distribution with a mean of 17.6 collinear genes pairs (2.25 mean-log) and a standard deviation based on the Ks_WGD_ being simulated.

**Figure 3:**
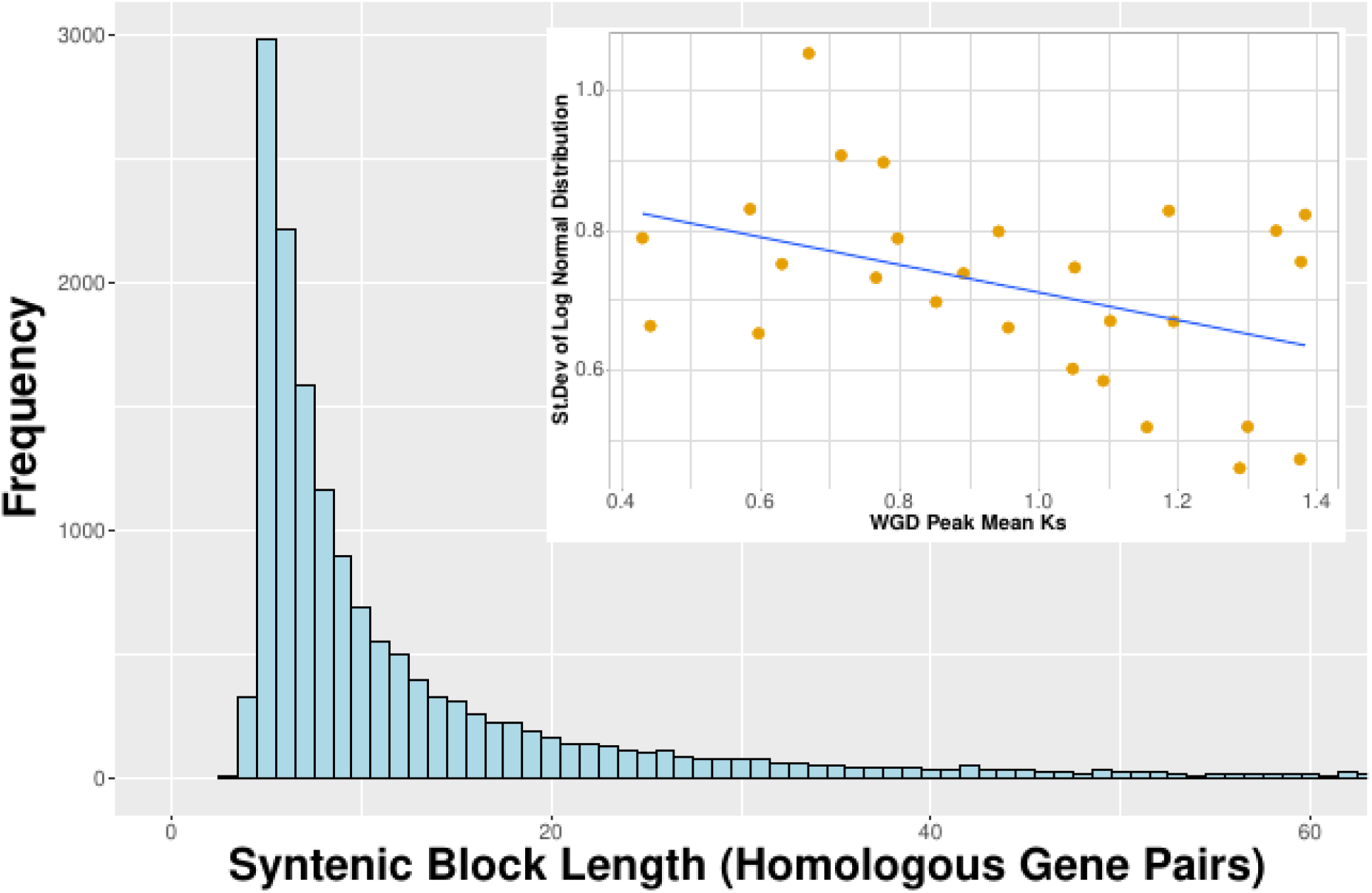
Comparison of the distribution of syntenic block lengths in terms of gene pairs across the entire dataset of 27 species against the distributions within each individual species. The histogram is syntenic block lengths in terms of gene pairs for the entire dataset of 27 species. The inset plots the standard deviation of log normal distributions fit to the syntenic block length distributions of each species against Ks_WGD_ of the youngest WGD in each. A linear model was fit to the data as described in section 3.1. Mean Ks_WGD_ was a significant predictor of the standard deviation of log normal distributions describing the distribution of block lengths in terms of gene pairs (F_1,27_= −2.242, p = 0.034, R^2^ = 0.162).

As syntenic blocks become broken up through time, the surviving collinear homologs also accumulate substitutions ***(29, 34, 43)***. Using the sequence divergence of these homologs it is possible to screen which syntenic blocks originate from different duplication events, such as the WGD in question or older WGDs and small scale duplications ***(29, 34, 43)***. Similar to the use of Ks plots to visualize the age distribution of paralogs in a genome, the mean synonymous divergence (Ks) of the collinear homologous gene pairs (Ks_block_) in each syntenic block pair can be plotted in a frequency distribution. As we could not *a priori* know the exact WGD each block originates from in our empirical data, we used EMMIX to fit a series of mixture models to a distribution of duplication node Ks values in gene family clusters calculated using DupPipe for each of the 27 species ***(73, 74)***. We then selected the model that fit the distribution for each species best based on the Bayesian Information Criterion (BIC) score and used Ks_pair_ with >50% likelihood assignment to the corresponding duplication peak to estimate a Ks boundary for each WGD peak. Syntenic blocks with a Ks_block_ within the bounds of these estimates were considered to have originated from the WGD. We then calculated the standard deviation of Ks_block_ values for each WGD peak and plotted it against Ks_WGD_ of each WGD. We fit a linear model to the data and found that Ks_WGD_ was a significant predictor of the Ks_block_ standard deviation (Fig. 4)(F_1,48_= 11.490, p < 0.001, R^2^ = 0.733). Using this model we then simulated the distribution of synonymous divergence among syntenic blocks based on the mean level of divergence for the WGD.

**Figure 4:**
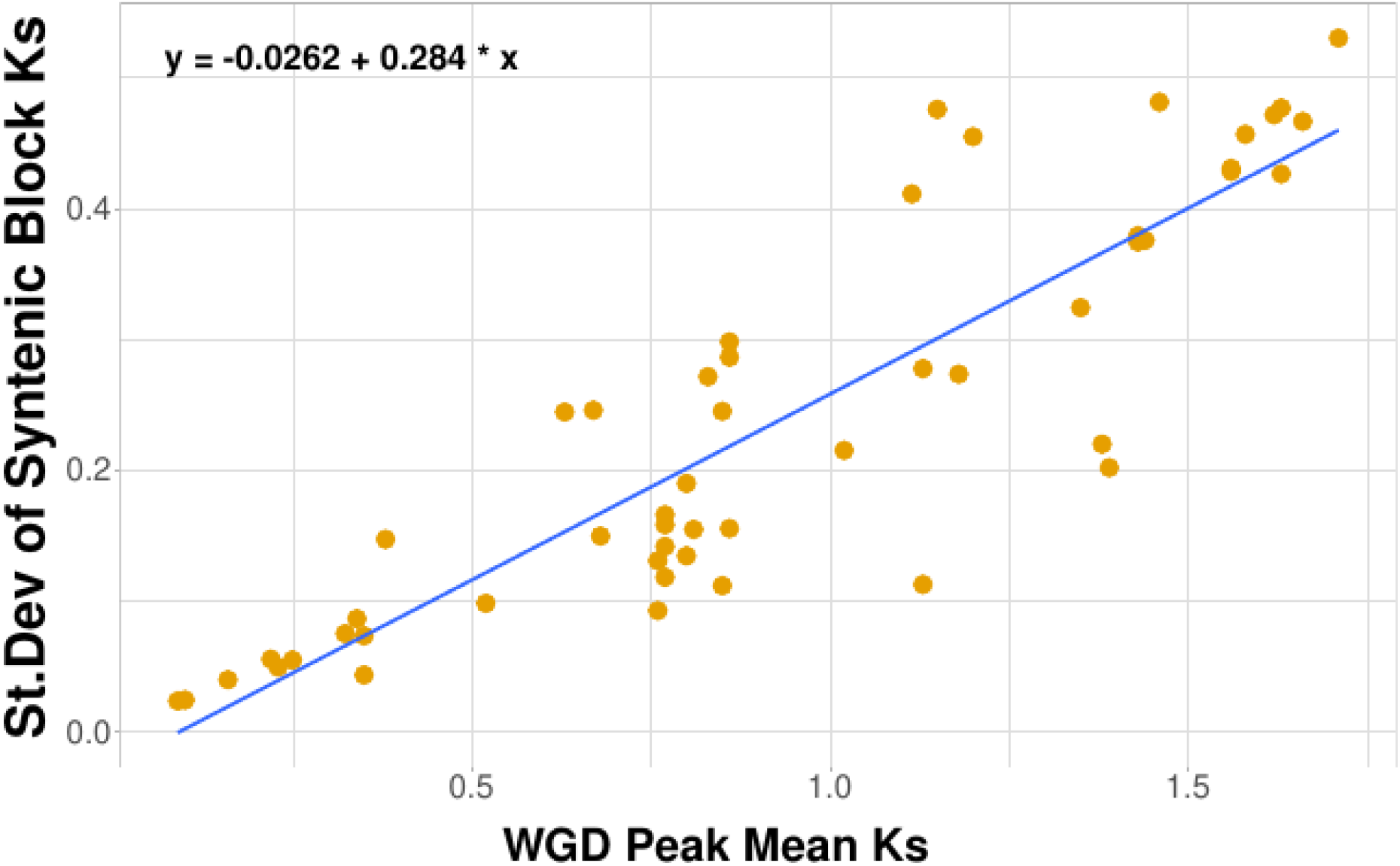
The standard deviation of the Ks_block_ for syntenic blocks plotted against the Ks_WGD_ for each WGD detected in all of the 27 species. A linear model was fit to the data as described in section 3.1. The Ks_WGD_ was a significant predictor of the standard deviation of Ks_WGD_ (F_1,48_= 11.490, p < 0.001, R^2^ = 0.733).

Not only do WGD differ in the variation of synonymous divergence among block pairs, but pairs of collinear homologs within those blocks can also accumulate different levels of synonymous divergence ***(34, 43, 75)***. To account for this variation, we calculated the standard deviation of Ks_pair_ within block pairs and plotted them against the Ks_block_ of each block. We fit a linear model controlling for block length and found that the standard deviation of Ks_pair_ within syntenic block pairs was significantly predicted by Ks_block_ (Fig. 5)(F_1,15,798_= 109.557, p < 0.001, R^2^ = 0.474). Using this model we simulated the level of synonymous divergence in collinear homologous gene pairs given the Ks_block_ of the syntenic block pair they are within.

**Figure 5:**
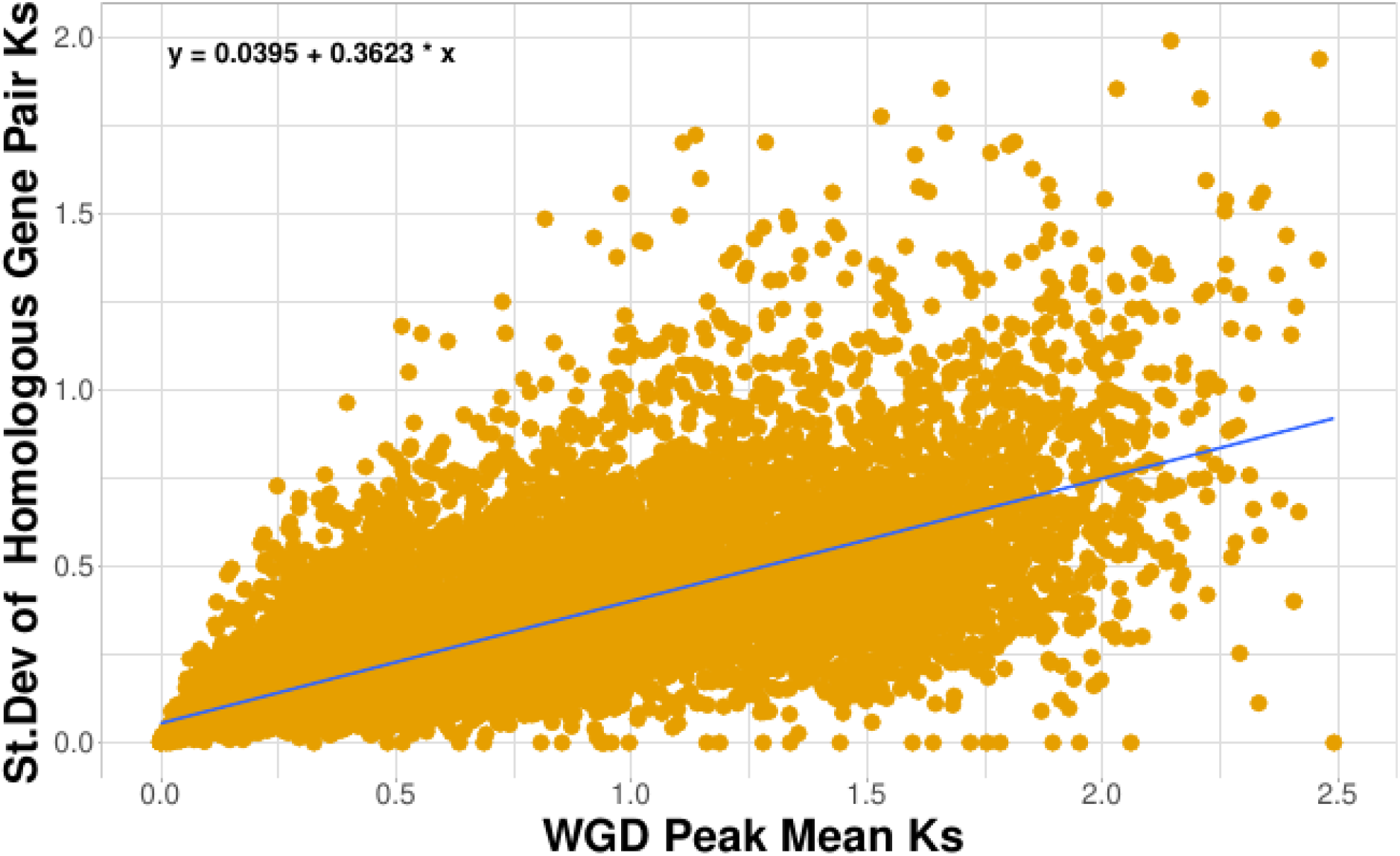
The standard deviation of the Ks_pair_ plotted against the Ks_block_ of each syntenic block pair detected in all of the 27 species. A linear model was fit to the data as described in section 3.1. The Ks_block_ was a significant predictor of the standard deviation of Ks_pair_ within each block (F_1,15,798_= 109.557, p < 0.001, R^2^ = 0.474).

### 3.2 Simulation

Using the models and distributions estimated from our 27 species, we simulated syntenic blocks from a variety of duplication histories in a format similar to the output of MCScanX (Fig. 1). This simulated training data was then used to train Frackify to classify paleologs in syntenic blocks detected by MCScanX or CoGe SynMap. The simplest of these training data simulated only a single WGD event. In single WGD simulations, we began by specifying the Ks_WGD_ and producing a list of featureless syntenic block pairs (Fig. 6A.1). The number of these block pairs can be determined by the user, allowing for flexibility in the scale of the simulation. Each block pair within the initial list was a different length randomly selected from a log normal distribution described by Ks_WGD_ and the model of block lengths for our 27 species (Fig. 3). We then simulated synonymous divergence in each block pair by assigning a random Ks_block_ from a normal distribution with a mean equal to the Ks_WGD_ and standard deviation specified by our model of Ks_block_ in section 3.1 (Fig. 4, Fig. 6A.2). Finally we incorporated the synonymous divergence in each collinear homologous gene pair within each block pair based on the Ks_block_ (Fig. 6A.3). This was done by selecting from a normal distribution with a mean equal to Ks_block_ and standard deviation from the model of Ks_pair_ within blocks from our 27 species (Fig. 5). At the end of the three previous steps we had a list of syntenic blocks similar to those found in MCScanX results for an individual WGD. If additional WGDs are desired, the user can specify to repeat this process for each. Using this approach we were able to produce simulated syntenic blocks for a wide variety of duplication histories similar to empirical examples (Fig. 1).

**Figure 6:**
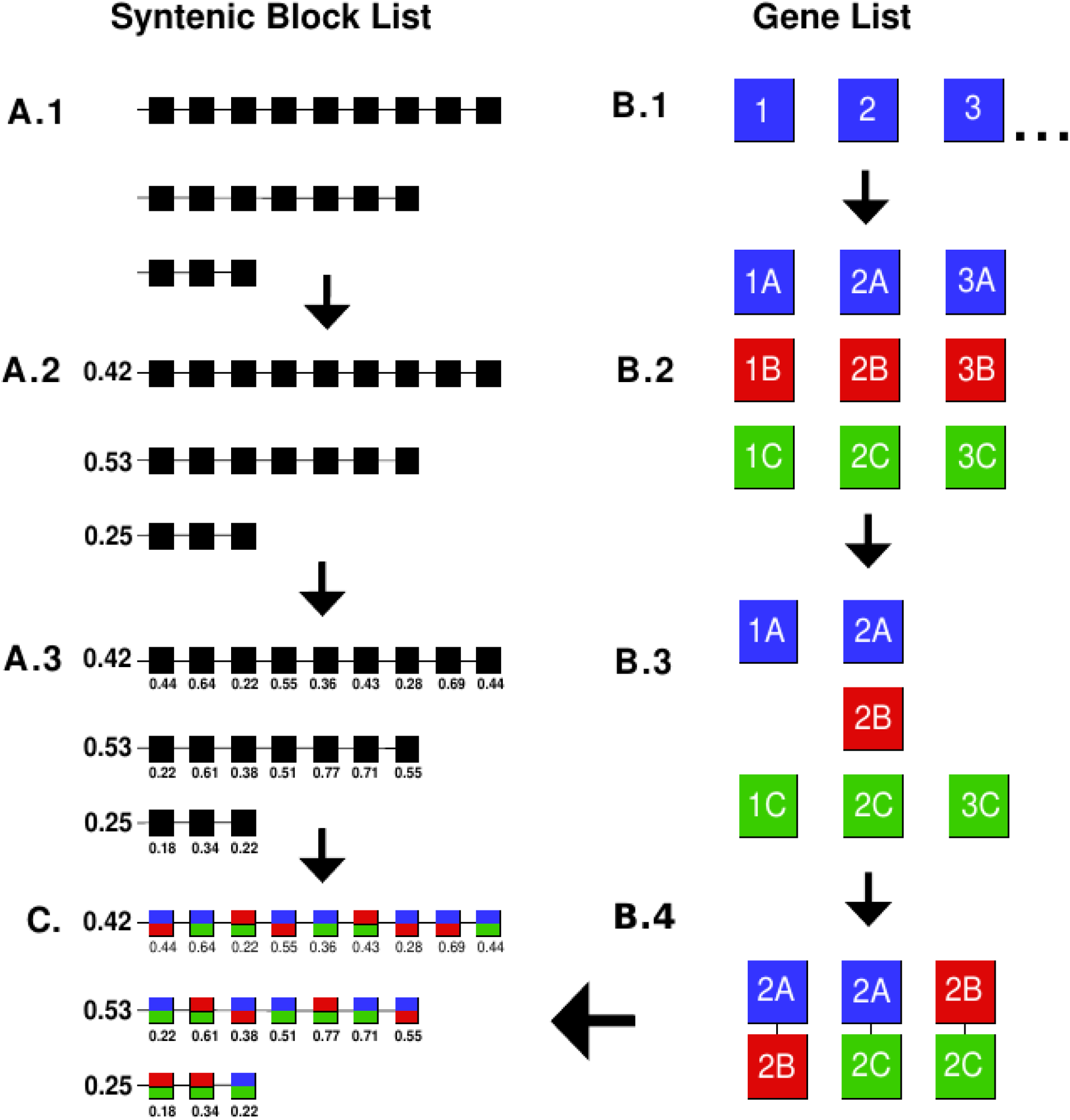
Flow chart for the basic steps used to simulate syntenic blocks. (A) Syntenic block simulation begins by (A.1) making a list of blocks, (A.2) assigning a Ks_block_ to each block, and (A.3) finally assigning individual Ks_pair_ to collinear homologous genes within each block. (B) To populate the syntenic blocks with genes, we first (B.1) generated a list of gene labels, (B.2) duplicated or triplicated the list depending on the nature of the polyploidy being simulated, (B.3) removed genes to simulate fractionation, and then (B.4) grouped the remaining genes into collinear homologous gene pairs. (C) Finally, the two lists were merged by randomly assigning homologous gene pairs to the appropriate syntenic blocks for the WGD of origin.

Within syntenic blocks produced by WGDs there are many classes of genes produced by different patterns of retention and additional small scale duplications ***(12, 23, 33)***. Pairs of homologous genes on syntenic blocks can represent paleologs retained as singletons, duplicate or triplicates, and non-paleologs produced by tandem duplication or gene conversion ***(11, 69, 76)***. To train Frackify to identify these classes of homologous gene pairs, we populated our simulated syntenic blocks with genes that reflect these different classes of gene duplications (Fig. 6B). In order to produce genes with varying duplication histories we began by producing a list of unduplicated genes. The length of this list was based on the number of empty collinear gene pairs in the simulated syntenic blocks from the most divergent WGD so that post-WGD and fractionation we would have enough syntenic WGD derived genes — syntelogs — to populate each simulated syntenic block (Fig. 6B.1). We then produced WGD derived syntelogs by duplicating or triplicating each gene in the list depending on if a paleohexaploidy or paleotetraploidy was being simulated (Fig. 6B.2)***(77)***. Next we simulated fractionation of these genes post-WGD by selecting a random number of genes to remove from each syntelog set between 0 and 3 (triplication) or 0 and 2 (duplication) (Fig. 6B.3). This resulted in approximately equal numbers of each paleolog class so that Frackify would be trained with a large sample of each. If a more complex duplication history was desired we duplicated and fractionated the gene list again to simulate repeated duplication events in the genome. Small scale duplications can also occur in genomes and result in genes which do not originate from our focal WGD ***(23, 33, 34, 69, 78)***. Genes from these additional duplications were simulated by producing a separate list of non-paleologs with a length equal to an arbitrary 15% of the paleolog gene list.

To produce the final simulated synteny dataset we merged our simulated list of paleologs and non-paleologs into syntenic blocks appropriate for each class (Fig. 6B.4). In the case of double retained paleologs, we expected each copy to be in WGD derived regions A and B that together make a single syntenic block pair ***(11, 34, 79)***. However, triple retained genes would be found in three regions (A,B and C), all being syntenic with each other. This resulted in pairwise block comparisons among all regions (e.g., A-B, A-C, and B-C)***(11, 38, 80)***. Most single retained genes would not be detected in self-self syntenic block pairs due to having lost their homologs, however a small number may be aligned with non-paleologs due to tandem duplications or gene conversion ***(38, 69, 76, 81, 82)***. To simulate this *a priori,* 10% of the single retained genes were randomly selected and paired with a non-paleolog to form a misaligned homologous pair. Once all pairs of collinear homologous genes were produced, they were randomly assigned to syntenic blocks from the WGD of origin to produce the final simulated self-self syntenic dataset (Fig. 6C).

Although self-self synteny data is useful for detecting WGD-derived syntelog sets, relying on it solely could lead to single retained genes being misclassified because they have lost their other paralogs ***(38, 81, 82)***. Previous methods have often identified single retained genes using gene tree information or syntenic comparisons to an outgroup ***(11, 38, 44, 81, 83)***. For Frackify we identified single retained genes using syntenic information from a related outgroup that the focal species diverged from prior to experiencing the WGD of interest. Similar to WGD, this speciation event can be detected in Ks plots due to a large peak of orthologous genes ***(28, 84–87)***. Genes in the focal species that were not already recognized as paleologs and were orthologous to an outgroup gene were considered to be single retained paleologs. This is because genes that have an ortholog in a pre-duplication outgroup—identified by both synteny and divergence—were likely present in the genome prior to WGD and the other copies were presumably lost during fractionation ***(83, 88)***. To simulate orthology with an outgroup we first simulated the shared duplication history prior to speciation by duplicating our self-self comparison list and removing syntenic blocks from the youngest WGD. Natural variation in synonymous divergence of the shared WGD in the outgroup and ingroup species was simulated by assigning new Ks_pair_ to each syntenic homolog pair using the same methods as the self-self comparison (Fig. 6A.2–3). The ortholog peak was then added by duplicating the gene list once more and assigning new Ks_pair_ to each ortholog pair with a mean Ks (Ks_orth_) between the focal and the second oldest Ks_WGD_. Although this final list represents our self-outgroup syntenic comparison, it is possible that some genes may be differentially lost between our focal species and the outgroup ***(11, 43, 89)***. To capture differential gene loss we randomly removed 10% of the genes from the outgroup, but users can specify different amounts of gene loss if desired.

Using our simulation framework we produced self-self and self-outgroup syntenic information for a variety of duplication histories. The simplest histories involved a single WGD event, which can have different levels of synonymous divergence. To represent these cases we produced 210 simulations with single WGD at Ks increments of 0.10 between 0 and 2 Ks. Some genomes have more complex duplication histories with multiple WGDs, a common occurrence in some land plant lineages ***(9, 13, 90, 91)***. For these complex duplication histories we also developed simulations with two and three WGDs. Similar to the single WGD simulations, we placed focal WGD at increments of 0.10 Ks starting at zero. For each Ks increment of the focal WGD we then simulated one or two additional WGD at distances in increments of 0.10 Ks away from the focal WGD until the most divergent WGD had a Ks of 2.0. Across the 1, 2, and 3 WGD scenarios we ran a total of 1540 simulations. To avoid producing an unnecessarily large amount of data we limited the scale of these simulations to produce an average of 10 syntenic block pairs per WGD compared to the average of 352 in the empirical dataset. These smaller scaled genomes contained an average of 634 genes per simulation. Across all 40,903 syntenic blocks we simulated a total of 976,896 paleologs and non-paleologs to train Frackify.

### 3.3 Training

The Frackify classifier was developed by training a boost decision tree using the XGBoost and Sklearn python libraries ***(61, 92)***. To prepare data for XGBoost, we parsed multiple features from the MCScanX like simulated data for each individual gene. The same feature extraction approach was also used for MCScanX empirical data. The first of these features included information pertaining to the duplication history of the organism or simulation, such as the Ks_WGD_ of each WGD and Ks_orth_ in the self-outgroup comparison. We also provided the difference in Ks between the focal WGD and possible neighboring WGD to help inform Frackify the possible degree of overlap in their Ks_block_ distributions. Next we focused on features of the genes themselves, beginning with self-self syntenic information. First we counted how many syntenic block pairs included the syntenic block each gene was found in. As some of these syntenic regions may have originated from older duplications, we then provided the number of syntenic block pairs which had Ks_block_ within three standard deviations of the focal Ks_WGD._. From these possible WGD derived syntenic blocks we counted the number of putative WGD derived syntelogs. As an example, we would expect a triple retained paleolog (gene A) to have a syntelog set size of three (A, B, and C) and be found in two syntenic block pairs (A-B and A-C). In the case of multiple overlapping WGD, it is possible this syntelog set may include genes that originated from an older duplication (gene O). To distinguish paleologs in cases like this, we then counted the total number of syntenic block pairs the syntelog set was involved in with Ks_block_ within 3 standard deviations of the Ks_WGD_. Using our triple retained paleolog example, if all syntelogs in the set are true paleologs we would expect three syntelogs (A, B, and C) and three syntenic block pairs (A-B, A-C, B-C). If one of the genes is from an older duplication (A, B, and O) we would have three syntelogs but only two syntenic block pairs (A-B, A-O). Finally, we averaged the Ks_block_ of each syntelog and reported how many standard deviations away from Ks_WGD_ the three closest syntelogs were. This initial set of summary statistics was chosen to summarize both the syntenic information for each individual gene and the syntenic relationships of its syntelogs.

In addition to features from the self-self syntenic data we also included self-outgroup syntenic information to both detect single retained genes and help parse out which genes in a syntelog set may originate from overlapping WGD. First we searched if each gene was found in syntenic blocks pairs with the outgroup at all. If a gene was in more than one syntenic block pair we then filtered out only the syntenic blocks with Ks_block_ within three standard deviations of the Ks_orth_. This process was repeated for all syntelogs in each set to gain a list of all the syntenic blocks that may be from the ortholog peak. We then filtered the outgroup genes in this list of syntenic blocks to find those that were orthologous to multiple syntelogs. If more than one orthologous outgroup gene was found, we calculated the mean Ks_block_ for all the syntenic block pairs that contained the outgroup genes. We selected the outgroup gene with the closest mean Ks_block_ to Ks_orth_ as it was most likely to be the ortholog of the WGD derived syntelog set. Finally, we listed the number of syntelogs in the set that matched the chosen orthologous outgroup gene. All of the features of the duplication history, self-self, and self-outgroup syntenic comparisons were formulated into a single row of a data frame along with the known class of the given gene.

With our data formatted we then trained our boosted decision tree while attempting to minimize overfitting ***(93, 94)***. First we assessed if the portion of the data used for testing and training could bias the model and lead to artificially inflated estimates of model accuracy while also searching for optimized hyperparameters using a Bayesian approach ***(95)***. This was done using a 5 k-cross fold validation, in which the data set is repeatedly split into training and testing data then an average accuracy of the model’s predictions is assessed. We repeated the 5 k-cross fold validation 200 times with different hyperparameters to minimize the negative mean absolute error ***(96–98)***. Once the final set of hyperparameters were selected we performed a final 5 k-cross fold validation, producing predictions for gene classes in the testing data with a mean accuracy of 96.05% varying by 2.32%. Given the low level of variance in how accurately the model identified gene classes, it is unlikely the model would be sensitive to the proportion of the data set that is used for training and testing. Based on this information we split our simulated dataset into 70% training and 30% testing data and trained the final model.

### 3.4 Testing and Validation

Using our simulated test data described above, we used a variety of standard approaches to evaluate the performance of Frackify’s predictions for the class of each gene. First we tested how well the thresholds for each class of gene were defined within the model using a Receiver Operating Characteristic curve (ROC)***(99, 100)***. For all classes, the rate at which false positives (FPR) was discovered remained low compared to the true positive discovery rate (TPR), suggesting the model was optimized correctly to avoid biasing towards one class of gene (Fig. 7). To evaluate the performance of Frackify further we made a confusion matrix which compared the classifications predicted by Frackify for each gene against the actual class, visualized as a heat map (Fig. 8)***(101)***. Across the 1,540 simulations Frackify predicted the majority (∼96–98%) of each classification category correctly (Fig. 8). Similarly, there was a low error rate between all classifications with no clear patterns of particular gene classes being mistaken for others. Between the ROC and heatmap, Frackify appears to perform well across a large set of simulated duplication histories, however it is possible certain scenarios are contributing the majority of the 2–4% of each class identified incorrectly.

**Figure 7:**
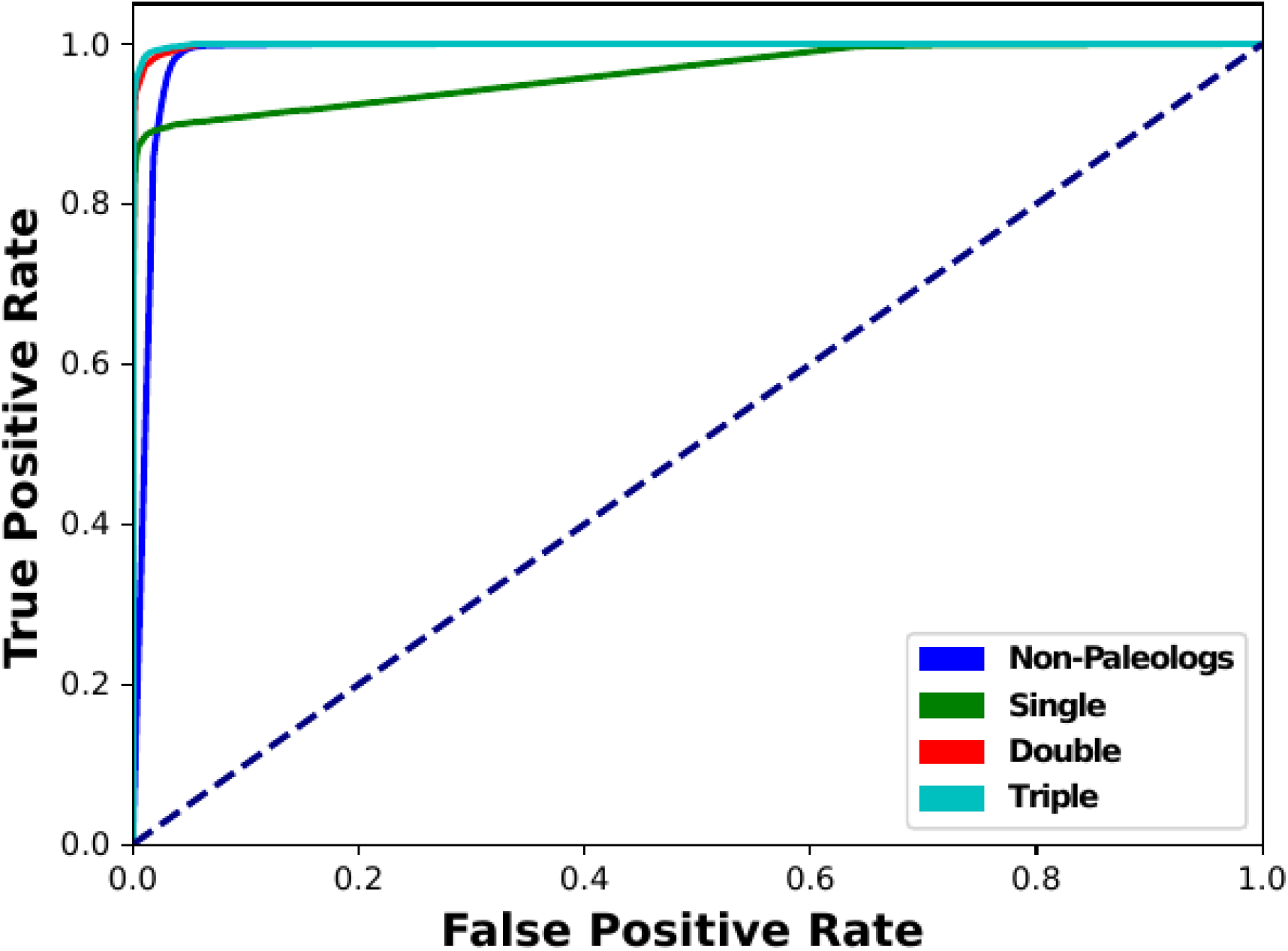
ROC curves for each of the paleolog classifications predicted by the trained model.

**Figure 8:**
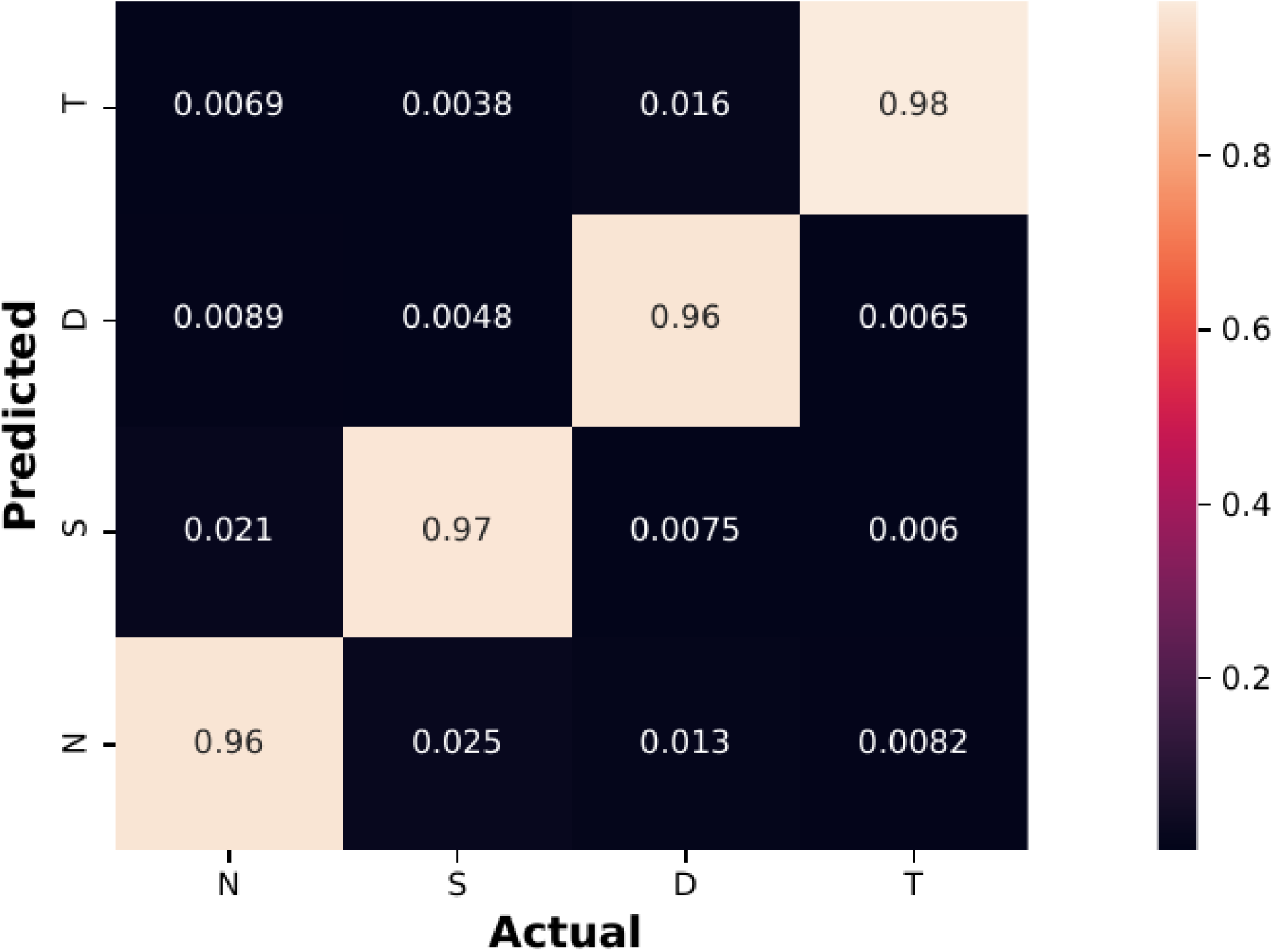
Heat map of the actual labels for genes in the testing data against the classes predicted by the trained model. N, S, D, and T represent non-paleologs, single, double, and triple retained paleologs respectively. The scale is the ratio of the genes in the actual class (columns) which were predicted by the trained model to be in each class (rows).

To further explore the small fraction of paleologs that were misclassified by Frackify, we considered several cases where paleolog classification might be challenging. For example, if multiple WGD peaks heavily overlap or speciation with the outgroup occurred soon after WGD ***(34, 83, 102)***. To evaluate Frackify’s performance in these cases we parsed out predictions for simulations that contained three WGD (Fig. 9). Despite the multiple overlapping WGD, non-paleologs were rarely misidentified as paleologs. However, Frackify less accurately identified different classes of paleologs in cases with highly divergent and overlapping WGD. There are several potential explanations for why this might be the case. As an example we considered the scenario where a given gene experiences a WGD, followed quickly by speciation, and then a second WGD. This scenario results in two sets of double retained paleologs from the younger WGD that are orthologous to two genes from the older WGD shared with the outgroup. In this case, Frackify does not misclassify these older duplicate genes as paleologs from the focal WGD. However, it may misidentify which ortholog each of the syntelogs matches, resulting in a triple retained paleolog set and a single retained paleolog. This issue could be compounded if one or more of the orthologs or paleologs are lost, such as in highly divergent WGD. Across most parameter space Frackify was able to discern these complex scenarios with a high degree of accuracy (∼85% or higher), but classification accuracy declined for Ks_WGD_ greater than 1.00 (Fig. 9).

**Figure 9:**
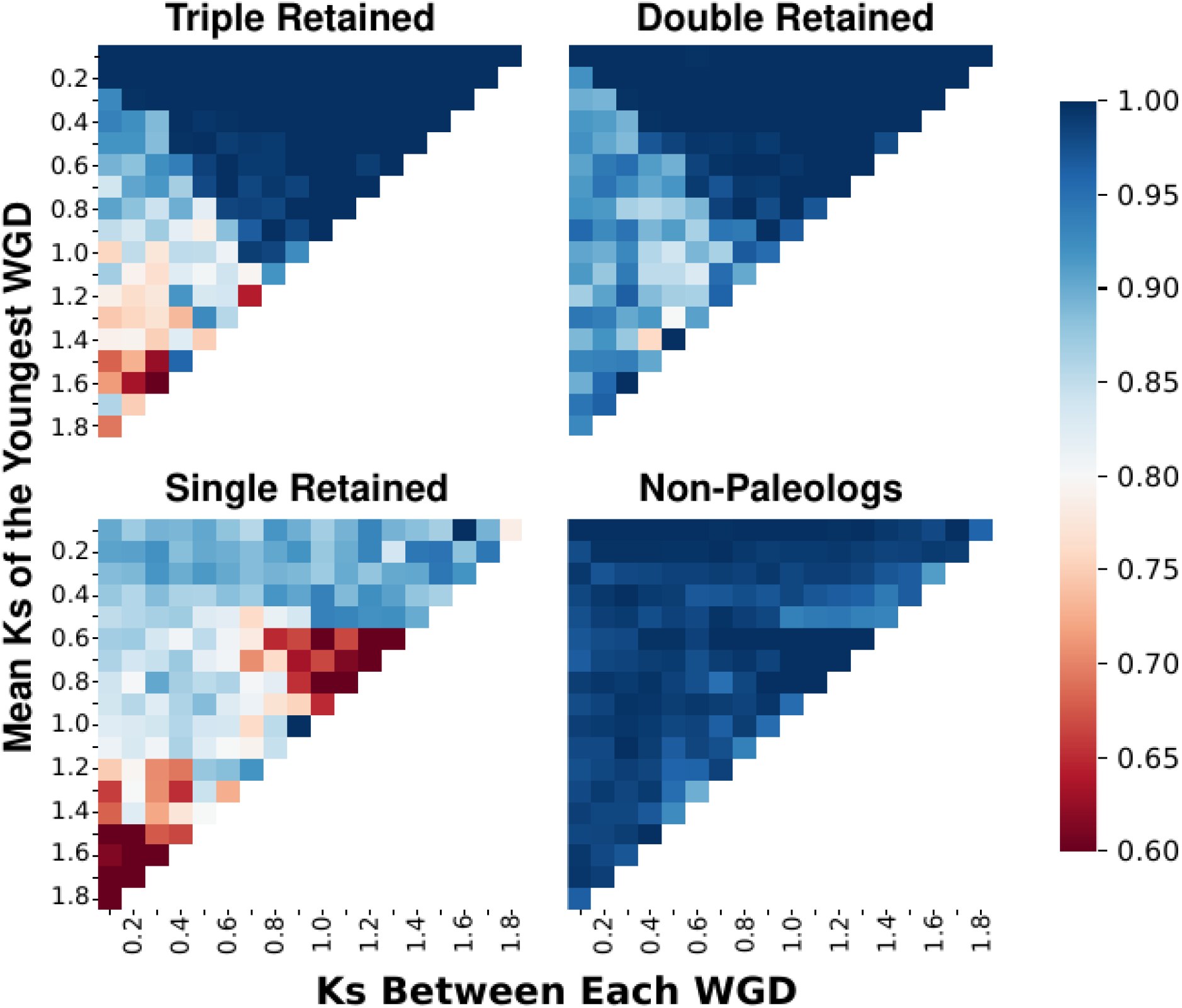
Heat maps for the accuracy of single, double, triple retained paleologs and non-paleologs predicted by Frackify across parameter space for simulations containing three WGD. The Y-axis represents the Ks_WGD_ for the youngest WGD, the X-axis represents the distance in Ks from the second and third oldest WGD to the focal WGD. Blank space represents parameter space where the oldest WGD occurs beyond the bounds of parameter space. The scale is the accuracy of predictions in terms of the ratio of correctly identified genes in each class out of all genes in that class.

To help understand why Frackify might sometimes misclassify paleologs, we explored the boosted decision tree itself. We used SHapley Additive exPlanation (SHAP) values ***(63, 103, 104)*** to help understand how XGBoost was making predictions for each class of gene. SHAP values are a useful data visualization tool to help interpret which features are having a significant impact on a machine learning model ***(103)***. In our case, we noted several important features that had consistently high values for several classes of genes (Fig. 10). In particular, we found that the number of syntelogs in a set and how many match to the same orthologous outgroup gene had high SHAP values across multiple gene classes. These features are likely major drivers of paleolog classification and in combination may explain why non-paleologs are rarely misclassified. For example, it is possible non-paleologs from more recent small scale duplications could be found in the focal WGD Ks peak, but they are not likely to be identified as paleologs because they do not have a clear ortholog in the outgroup. Similarly, these primary features may be used in combination with other features to help discern more complex scenarios. For example, if a set of paleologs contains a paralog that actually originates from an older, shared WGD, that paralog will be syntenic to multiple regions of the outgroup genome. These additional syntenic regions will result in an inflated number of putative orthologous outgroup genes, lending evidence that the paleolog set is contaminated with paralogs. These secondary features can be highly gene class specific, evident by their SHAP values. For example, the most important features for the classification of double retained paleologs was the number of standard deviations away from Ks_WGD_ the mean Ks_block_ of each syntelog (Fig. 10). These features are likely how the model discerns if syntenic block pairs in the self-self comparison originated from the focal or overlapping WGD. Although we do not know the exact way these features are used to make final predictions, their importance to the model is useful to understand how we may improve Frackify’s classifications in the future and for comparing Frackify to other approaches.

**Figure 10:**
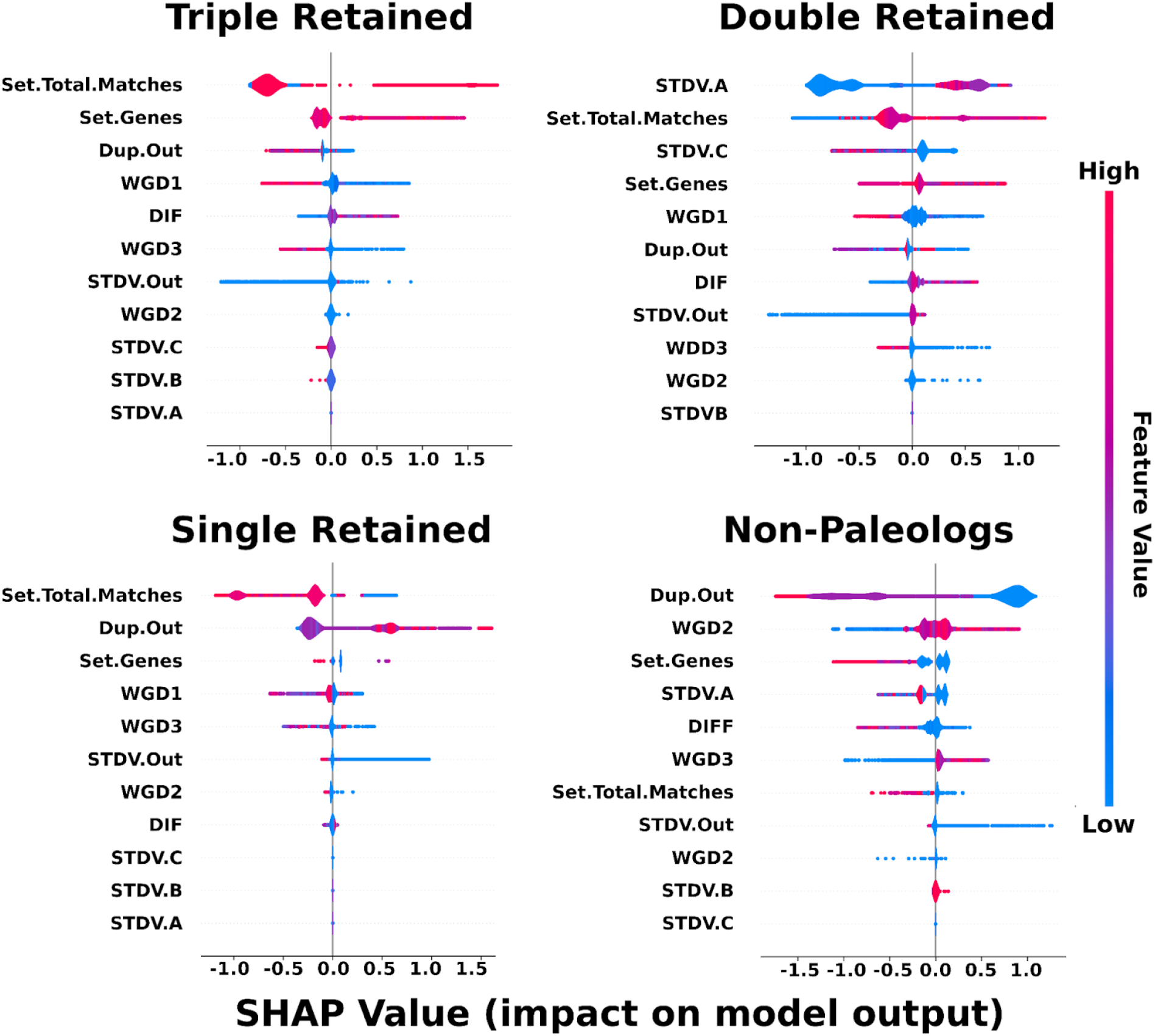
SHAP values for each class of genes predicted by Frackify. Violin plots represent the distribution of SHAP values for a given class of gene with outliers represented as dots. The color of the violin plot indicates the feature value for the model’s output, while the X-axis represents the SHAP value for a given point. Each feature describes the following properties; WGD (1–3) is the K_WGD_ of each WGD in the species, DIF is the difference in Ks between the two youngest WGD, Syn.Genes describes how many genes are in the WGD derived syntelog set, STDV(A–C) describes how many standard deviations away from the WGD each gene in the set is, Dup.Out represents if the a gene matches to an outgroup gene in the orthology peak, Syn.Matches described how many of the genes in the WGD derived slytelog set match to the outgroup gene, and STDV.Out is how many standard deviations away from the orthology peak the outgroup gene is.

### 3.5 Empirical Examples

Although Frackify performed well when validated with simulated test data, we wanted to test the model’s performance on empirical data sets. To accomplish this we compared Frackify’s classifications with paleolog inferences from POInT for six plant species ***(11, 14, 80)***. Many of these species are of agronomic importance and have experienced multiple WGDs, making them a useful testing ground for paleolog inference tools ***(105–108)***. In *Thellungiella halophila*, *Capsella rubella*, and *Eutrema parvulum* we analyzed the *At*-α duplication event, estimated to have occurred 14.5–86 Ma with a median WGD peak Ks near 0.70 ***(3, 66)***. For *Brassica rapa*, *B. oleracea*, and *Crambe hispanica* we analyzed the Brassiceae whole-genome triplication, with a median WGD peak Ks near 0.30 ***(3, 14)***. Similar to POInT, we used *Cleome violacea* and *Arabidopsis thaliana* as outgroups for the Brassiceae whole-genome triplication and *At*-α WGD respectively ***(11)***. We compared the POInT paleolog inferences in these six genomes to the less rigorous Ks + Synteny overlap based approach discussed in the introduction and new classifications from Frackify.

Across all six species we found that Frackify identified genes as paleologs with a large degree of overlap with other approaches, but also identified many additional paleologs. We visualized this overlap using euler plots from the eulerr R library (Fig. 11)***(109)***. Over half of the paleologs identified were shared across more than one method (58.7% total, 53.0–62.1% across species)(Fig. 11). A relatively small proportion of unique paleologs classifications came from POInT and the Ks + Synteny overlap approach (5.1% total, 0.3–5.5% across species)(Fig. 11). Similarly, Frackify identified 95.3% and 92.1% of the paleologs identified by Ks + Synteny and POInT respectively. In the case of POInT, Frackify also maintained a high degree of agreement in terms of the retention status of each paleolog classification (81% average, 63 – 89% range)(Fig. 12). These results suggest that Frackify is not only able to identify the majority of paleologs previously identified by other methods, but also their retention status. In addition to these findings, a significant proportion of paleologs identified were unique to Frackify (36.2% total, 33.0 – 39.0% across species)(Fig. 11).

**Figure 11:**
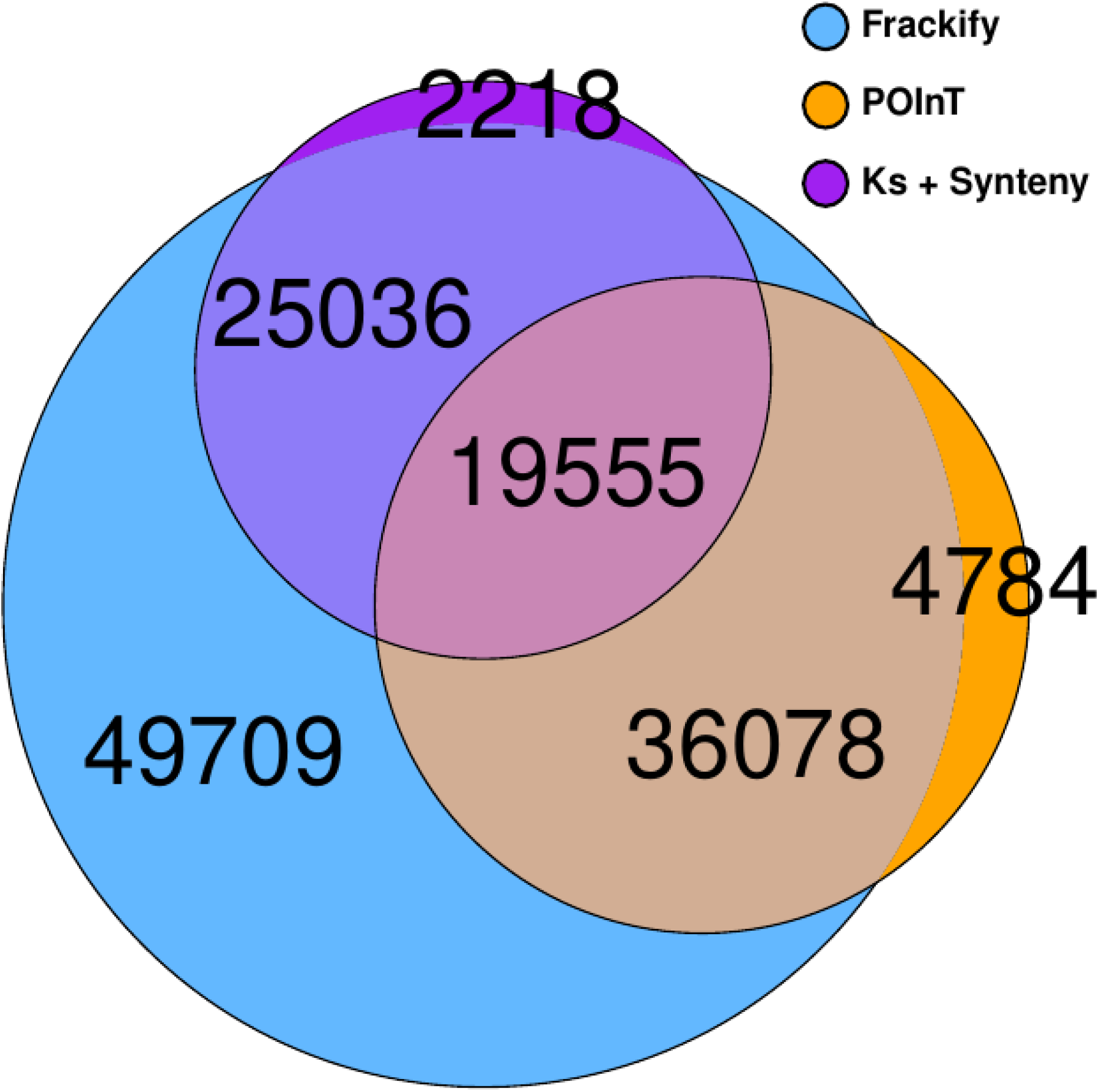
Euler plot for paleolog identifications across all 6 species used in the empirical data set comparison. The area of each region is approximately proportional to the percentage of the total dataset it represents.

**Figure 12:**
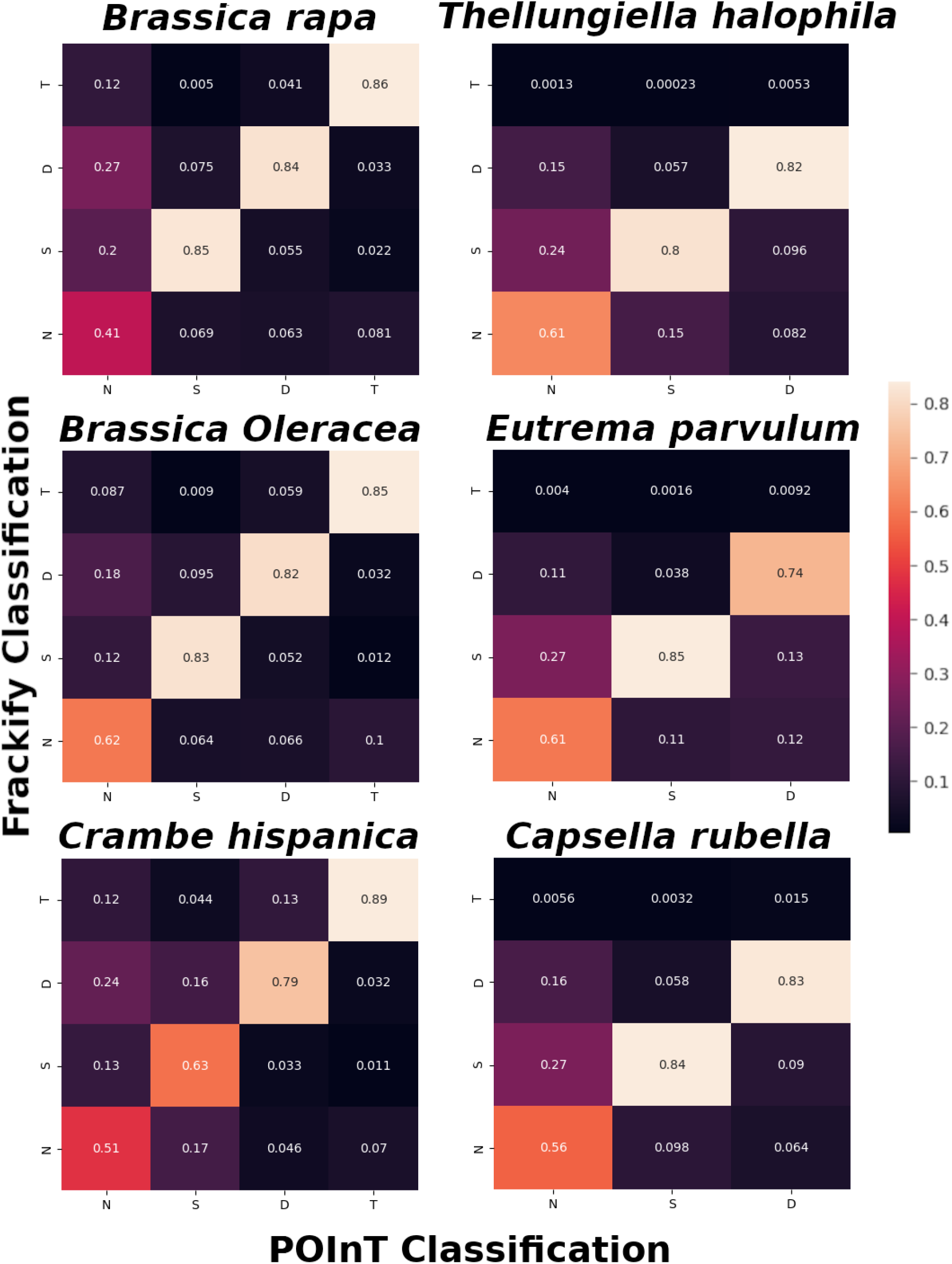
Heat maps for predicted class of each gene made by Frackify compared against inferences made by POInT in the six species. On the left hand column are paleohexaploids with up to three gene copies retained, while the right hand column contains paleotetraploids that only retain up to two copies of each gene. Color indicates the level of agreement between the two methods scaled as the ratio of the total number of genes in each class inferred by POInT (Columns) compared to Frackify’s predictions (Rows).

To further analyze the additional genes identified by Frackify and qualitatively assess if they are generally accurate or false positives, we conducted a more detailed comparison of classification differences between POInT and Frackify that leveraged SHAP values from the model. For all classes of paleologs identified by POInT we noted a high degree of agreement with Frackify, suggesting the model was able to recognize these classifications and make accurate predictions (81% average, 63 – 89% range)(Fig. 12). The major source of disagreement between Frackify and POInT occurred in non-paleologs. It is possible these non-paleologs were simply not analyzed by POInT due to being in DCS blocks not conserved across all species ***(11, 44)***. This was supported by the observation that 46,195 (62%) of the additional paleologs identified by Frackify were not found in the output files of POInT. To assess if these additional paleologs were false positives, we used SHAP values to compare their features. The most consistently important features for paleolog classification in Frackify were 1) the WGD derived syntelog set size and 2) how many genes in the set matched to the same outgroup gene in the ortholog peak (Fig. 10). It is possible that some of the genes identified using these metrics may originate from other WGDs in the genomes. To test if this was the case, we examined how many paleologs were found in any syntenic blocks with a mean Ks within three standard deviations of Ks_WGD_ and Ks_orth_. Filtering by this metric we found that out of 74,745 total unique paleologs Frackify identified across all six species, only 389 were found outside our expected Ks_WGD_ and Ks_orth_ distributions. Although some of these paleologs (14,453) were also located in syntenic blocks pairs originating from older WGD, our analyses suggest that they were duplicated in the youngest WGDs rather than being false positive classifications. Thus, many of the paleologs uniquely recovered by Frackify are not clearly false positives and are likely paleologs missed by other methods.

### 3.6 Summary

Frackify demonstrates how machine learning approaches can be used to explore the evolution of complex, duplicated genomes. By combining multiple features of genome synteny and divergence, Frackify is able to relatively rapidly identify paleologs in diploidized species with both simple and complex duplication histories. Frackify uses a gradient boosted decision tree trained with simulated syntenic blocks developed from the empirical data of 27 paleopolyploid plant species. In testing, Frackify performed well under a wider variety of duplication scenarios, including those with multiple, overlapping WGDs. Finally, we compared Frackify against other sophisticated methods used for identifying paleologs. Across six empirical datasets, Frackify maintained a high degree of consistency with methods such as POInT while also identifying a large set of additional putative paleologs in regions of the genomes not analyzed by these methods. With this tool, researchers can quickly identify paleologs in species of interest while also gathering information about patterns of paleolog retention.

Using SHAP values we were also able to gain a deeper understanding of the importance of different genomic features for identifying paleologs. Previous methods have often relied on syntenic information or synonymous divergence and applied complex models or heuristic filters to identify paleologs ***(11, 27, 28, 30, 32–36, 80)***. SHAP values uncovered that syntenic information was the primary feature used by Frackify to classify paleologs. However, Ks information was critical for assigning syntenic blocks to particular WGDs or other duplication events (Fig. 10). Similarly, methods which utilize an outgroup, such as POInT, Frackify, or Quota-Align, are able to resolve paleolog classifications in regions of the genome that are not necessarily detected during self-self syntenic comparisons ***(11, 34, 44, 80)***. As a result, carefully choosing an outgroup to compare against is critical for marking accurate inferences. These inferences suggest that improving ortholog identification with outgroup genomes could be important for better classification and characterization of paleologs. Further merging of synteny and gene family phylogenetic data should improve current paleolog classifications while also allowing for multi-species comparisons. Approaches such as Frackify provide one possible path to combining these disparate types of data to make genomic inferences.

### 3.7 Using Frackify

Frackify has several folders that the user will need to populate with data before running the program. When populating these folders users should use the same gene names for each file and make the prefix (Example: “species”) of the file consistent. Examples have been placed in the Git repository for users to emulate formatting.

1. CDS: This folder contains CDS fasta files for each diploid species the user intends to run Frackify on. Use the file naming format “species”.cds
2. Self-Self: For each species place either a CoGe Synmap or MCScanX output file for the self - self synteny comparison. These files should have Ka/Ks values appended to them and the gene names match the CDS file. Use the file naming format “species”-Self.txt
3. Self-Out: This folder will contain the same files as described for Self-Self, however for a self - outgroup syntenic comparison. This outgroup should be a diploid closely related to the focal species that does not share the WGD you are analyzing for paleologs. Use the file naming format “species”-Out.txt
4. In the main folder: Supply a CSV file containing information about each species you intend to run Frackify on. This file will have 5 columns: 1) the first will contain each species name; please make this the same as your file names. 2) The second column will contain the Ks_orth_ of the outgroup, typically found between the focal and next oldest WGD. 3–5) The third through fifth columns will contain the Ks_WGD_ of the three youngest WGD in the species in ascending order. If only one or two WGD are present in the genome, list the others as “0”. An example of this format can be found in the Frackify git repository (https://gitlab.com/barker-lab/frackify/-/tree/master).

The first script to run in the Frackify pipeline will produce an intermediate file that contains the features of each gene summarized in a table. This can be produced using the following command.

> *python3 Survey.py “species_info_.csv”*

Following the completion of the script two files will be generated in the “*Translated_Data*” folder. The first will be the list of features for each gene (*“species.Forest.csv”)* the second will contain the list of all syntelog sets from the self-self and self-outgroup comparisons (*“species.Groups.csv”*). Users can then produce predictions using the main Frackify command.

*python3 Frackify.py “Translated_Data/species.Forest.csv”*

The Frackify script will produce a final csv file in the Predictions folder that contains a list of all the genes in the CDS file and their classification. For single, double, and triple retained genes a separate file will be produced that contains the WGD derived syntelog sets and the outgroup gene they match in each row.

Alternatively, users can train a new boosted decision tree using different simulations. Simulations can be produced by supplying a CSV file with a list of each desired scenario. Users will provide unique names for each scenario to be simulated in the first column. In the second column, provide the number of syntenic blocks to be simulated per WGD. The third column allows the user to specify additional fractionation from 0–1 where 0 is no additional fractionation and 1 would represent a complete loss of all paleologs. In the fourth column the user can select which WGD will be used for paleolog simulation (if there is more than one WGD in the simulation). In column 5 specify the Ks_orth_ of the outgroup ortholog divergence. Columns 6–8 should contain the Ks_WGD_ of up to three WGD in order of ascending divergence. If less than three WGD are specified, place zeros in the remaining WGD Ks columns. The final three columns are used to identify if each WGD is a duplication (2 = paleotetraploid) or triplication (3 = paleohexaploidy). Similar to Frackify, running these simulations requires only one command.

> *python3 Sim_Stations.py “file_of_simulation_parameters.csv”*

Running the Sim_Stations.py script will populate the Simulated_Data folder with data from the simulations that can then be used to train new gradient boosted models. The training script “*Fracking.Rig.py*” has two commands, one that will generate Bayesian optimized hyperparameters for this training set ***(1)***, and another that will add a new trained model in the “*Models*” folder ***(2)***. If users wish to use their new model in Frackify, please alter the “Frackify.py” script at line 25 to load the new model.

> *python3 Frack.Rig.py “1 or 2”*

## 4. Notes

1. Frackify depends on MCScanX and SynMap results to make inferences, and as a result it is inherently limited by the quality of syntenic information identified using such tools. Users should consider this when interpreting results as syntenic blocks can be missed in the case of poor genome assembly quality, when using a highly divergent outgroup, or altering MCScanX/SynMap.pl settings. Below we have included several ways to help determine if paleologs are being missed due to syntenic block inferences.

a. In the final step of Frackify, genes predicted as paleologs are grouped together using the self-outgroup and self-self syntenic information. During this process it is possible that some syntelogs in a given set have disagreeing predictions; for example a triple retained group being split into a single retained gene and double retained pair. Frackify will print out the number of genes that suffer from these problems, a large amount suggesting some syntenic blocks are being undetected.
b. The same issue described in point “a” can be observed within the “Groups” file. This problem will manifest as some outgroup genes occurring more than once in the file, suggesting the self-self comparison is missing a large number of syntenic blocks.
c. If more than 10% of genes display either of the issues in points “a” or “b” we recommend users refer back to their MCScanX or SynMap results to diagnose why syntenic blocks are being undetected.
d. The simplest and most influential method to help limit the number of disagreeing paleolog classifications is to be careful when selecting an out group. Users should take time to carefully identify the least divergent out group possible while also prioritizing genome quality. Highly divergent genomes or those with poor assemblies can lead to inflated false positives and false negatives due to the limitations of synteny detection tools.
2. In the case where older WGD in the outgroup overlap with the orthology peak, it is possible to use quota-align in SynMap to help identify orthologous blocks if the syntenic depth of the WGD is known *a priori* ***(34)***. While this can limit false positives, it can also result in false negatives for single retained paleologs.

